# Variable Effects on Growth and Defence Traits for Plant Ecotypic Differentiation and Phenotypic Plasticity along Elevation Gradients

**DOI:** 10.1101/435453

**Authors:** Moe Bakhtiari, Ludovico Formenti, Veronica Caggía, Gaëtan Glauser, Sergio Rasmann

## Abstract

Along ecological gradients, ecotypes generally evolve as the result of local adaptation to a specific environment to maximize organisms’ fitness. Alongside ecotypic differentiation, phenotypic plasticity, as the ability of a single genotype to produce different phenotypes under different environmental conditions, can also evolve for favouring increased organisms’ performance in different environments. Currently, there is a lack in our understanding of how varying habitats may contribute to the differential contribution of ecotypic differentiation and plasticity in growth versus defence traits. Using reciprocal transplant-common gardens along steep elevation gradients, we evaluated patterns of ecotypic differentiation and phenotypic plasticity of two coexisting but unrelated plant species, *Cardamine pratensis* and *Plantago major*. For both species, we observed ecotypic differentiation accompanied by plasticity in growth related traits. Plants grew faster and produced more biomass when placed at low elevation. In contrast, we observed fixed ecotypic differentiation for defence and resistance traits. Generally, low elevation ecotypes produced higher chemical defences regardless of the growing elevation. Yet, some plasticity was observed for specific compounds, such as indole glucosinolates. We speculate that ecotypic differentiation in defence traits is maintained by costs of chemical defence production, while plasticity in growth traits is regulated by temperature driven growth response maximization.

## Introduction

Species with extensive geographical ranges tend to exhibit large intraspecific variation in most functional and phenotypic traits. Such geographic variation can lead to the evolution of morphologically and functionally different genotypes or ecotypes (Hufford and Mazer, 2003; Kawecki and Ebert, 2004; Savolainen *et al.*, 2007). Ecotypes are comprised of genetically distinct population of a given species retaining traits that maximize fitness leading to local adaptation to particular local abiotic and biotic conditions (Kawecki and Ebert, 2004). Phenotypic variation within a species due to heritable ecotypic differentiation is further distinguished in different habitats by phenotypic plasticity. Phenotypic plasticity refers to the ability of a single genotype to produce different phenotypes under different environmental conditions. Plasticity itself can also be selected for and evolve differently for different developmental, physiological, and reproductive traits or in different habitats in order to optimize organisms’ performance (Bradshaw, 1965; Gotthard *et al.*, 1995; Lortie and Aarssen, 1996; Murren *et al.*, 2015; Scheiner, 1993; Sultan, 1987; Sultan, 2003). Species with greater adaptive plasticity respond more acutely to environmental changes, and may be better able to survive in novel environments allowing their rapid geographical spread inhabiting a broad range of environmental conditions (Baker, 1974; Oliva *et al.*, 1993; Spencer *et al.*, 1994), thus promoting local adaptation (Baldwin, 1896; Ghalambor *et al.*, 2007; Price *et al.*, 2003).

As sessile organisms, plants should experience strong local adaptation to local climate that strongly affects plants’ fitness. For instance, with temperature transitions across species’ latitudinal ranges or altitudinal niches- spanning low to high elevations-, plants tend to evolve to produce smaller seeds, to have earlier phenology, slower growth rates, and display greater investment in clonal reproduction (e.g. Chapin and Chapin, 1981; Körner, 2003; Moles *et al.*, 2007; Montague *et al.*, 2008; Pilon *et al.*, 2003). At the community level, the emergence of interspecific interaction clines out of biogeographical clines is also expected. Since the initial Dobzhansky’s postulate of a potential correlation between biotic interaction strength and trait values for traits mediating such interactions (Dobzhansky, 1950), a large volume of literature has focused on plant-herbivore interaction (Bolser and Hay, 1996; Coley and Aide, 1991; Schemske *et al.*, 2009). More specifically, it is expected that increased herbivory pressure in the tropics should favour the evolution of more potent defences in plants (Coley and Barone, 1996; Moles *et al.*, 2011; Pellissier *et al.*, 2014; Pennings *et al.*, 2001; Rasmann and Agrawal, 2011; Siska *et al.*, 2002; Woods *et al.*, 2011). Furthermore, a decrease in species diversity at high altitude can also be associated to a reduction in species interaction, and in turn, a relaxation of plant defences across scales, such as at the community level (Callis-Duehl *et al.*, 2017; Descombes *et al.*, 2016; Kergunteuil *et al.*, 2018), at the interspecific level (Defossez *et al.*, 2018; Pellissier *et al.*, 2012), as well as at the intraspecific level (Pellissier *et al.*, 2014; Scheidel and Bruelheide, 2004; Zehnder *et al.*, 2009). In analogy with latitudinal gradients, elevation gradients are emerging as optimal tools for studying plant trait variation along ecological clines that occur over short geographic distances (Körner, 2007). Indeed, plant adaptation to habitat-specific abiotic and biotic factors can be studied along elevation transects regardless of biogeographic history, gene-flow barriers, and within homogenous macroclimatic conditions (Rasmann *et al.*, 2014; Sundqvist *et al.*, 2013). Along environmental gradients, trait-mediated local adaptation of plant ecotypes is the result of selection for fitness maximization given the local biotic and abiotic conditions. Within genetically determined trait differences between ecotypes, variation emerges from phenotypic plasticity if plasticity for such trait expression does not come with relative costs to fitness (Gratani *et al.*, 2003; Van Tienderen, 1989).

Plant growth and defence related traits have been shown to vary in response to different conditions. For instance, high and low elevation *Plantago lanceolata* ecotypes growing at two temperature regimes (15 and 25 °C) showed strong plasticity in growth (i.e. both genotypes grew similarly within each environment), while their resistance to generalist herbivores reflected genetically-fixed patterns; high-elevation ecotypes were always less resistant, independently of the temperature regimes (Pellissier *et al.*, 2014). Such differences in plasticity would suggest that ecotypes that, at high elevation, produce lower amounts of constitutive defences were favoured by selection, and growing in warmer temperatures could not modulate this pattern of defence production. Similar reciprocal transplant experiments have been classically used to measure the extent of ecotypic differentiation and phenotypic plasticity (Nahum *et al.*, 2008). The predictions being that ecotypes adapted to one environment should change their phenotypes when place in a novel environment given their genetic constraints. Coupling reciprocal transplant with common garden experiments is critical because phenotypic plasticity of growth and defence traits in response to growing conditions can also generate clines, and such plasticity can obscure genetically based trait expression.

With this study, we aimed at measuring the magnitude of ecotypic differentiation and plasticity in growth and defence traits of two unrelated plant species with similar geographical distribution along elevation gradients in the Alps (Supplementary Fig. S1). Specifically, we collected seeds of four populations of *Cardamine pratensis* (Brassicaceae) and six populations of *Plantago major* (Plantaginaceae); half of the populations were native to low elevation and the other half to high elevation. We grew high and low elevation ecotypes at both their native or non-native elevation range using two common gardens along a mountain transect, and we assessed variation in growth and defence (secondary metabolite) related traits. Based on the theoretical framework as shown in Fig. 1 (Leggett *et al.*, 2014; Schlichting and Pigliucci, 1998), we expected five contrasting scenarios: 1) to observe no ecotypic variation or plasticity when the traits remain constant across ecotypes and environments (Fig. 1A). 2) To observe ecotypic differentiation (ecotype effect only) with no plasticity when trait variation remains constant across elevations for a given ecotype but different ecotypes would exhibit different trait values (Fig. 1B). 3) To detect plasticity without ecotypic differentiation (elevation effect only) when both ecotypes show trait variations across different growing elevation, without significant difference between ecotypes (Fig. 1C). 3) To observe ecotypic effect accompanied by plasticity if different ecotypes exhibit differential values both from one another and at different growing elevation (elevation and ecotype effects) (Fig. 1D). Finally, we would expect to observe plasticity through genotype by environment effect when the interaction of ecotype and elevation explains the traits value (elevation × ecotype effect) (Fig. 1E). Overall, this study builds towards a better understanding of the ecological and evolutionary drivers of pathways mediating plant phenotypic variation in growth versus defence traits along ecological clines.

**Fig. 1.**
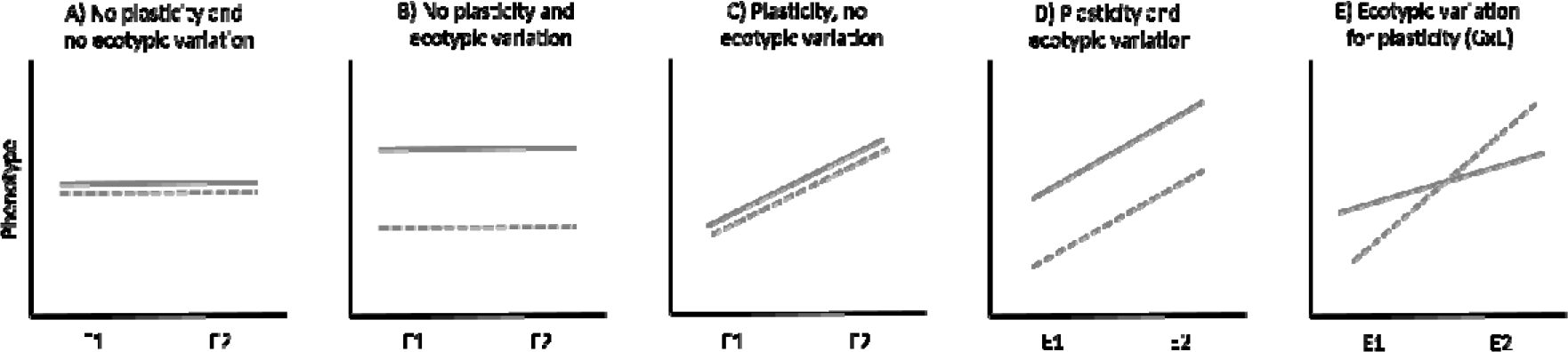
Theoretical framework for measuring ecotypic differentiation and phenotypic plasticity using reciprocal transplant experiments and reaction norms. The different panels represent all likely scenarios.

## Material and methods

### Plant materials

*Cardamine pratensis* is a rhizomatous perennial herb that grows in a variety of habitats including nutrient-rich meadows, pastures, and forests and is common throughout Europe and in Central and Eastern Asia (Hultén and Fries, 1986). *C. pratensis* populations cover a wide elevation range, from sea level to 1600 meters above sea level (Aeschimann *et al.*, 2004), and flowers from April to June. Flowers are self-incompatible, and plants generally produce clonal offspring as new rosettes, especially under moist conditions (Lövkvist, 1956), and are considered hemicryptophyte (i.e. a long-lived geophyte with overwintering green leaves). *Cardamine pratensis* contain glucosinolates (GLS), which, when in contact with myrosinases, enzymes present in separate compartments of the cells, are degraded into glucose and sulphate, along with various nitrile, isothiocyanate, and thiocyanate molecules that are toxic or deterrent to generalist insect herbivores and some pathogens (Giamoustaris and Mithen, 1995; Hopkins *et al.*, 1998; Kliebenstein *et al.*, 2002; Lambrix *et al.*, 2001). GLS are often classified into three classes of compounds depending on their side-chain: aliphatic, indole and aromatic, several of which have been shown to be effective against generalist and, to some extent, against specialist herbivores (Daxenbichler *et al.*, 1991; Louda and Rodman, 1983; Montaut and Bleeker, 2011). GLS are known to vary quantitatively and qualitatively (Kliebenstein *et al.*, 2001; Mauricio, 1998). In addition, phenotypic plasticity in GLS production has been previously observed in wild brassicaceous species (Agrawal *et al.*, 2002). For instance, GLS profiles of *Boechera stricta* were strongly plastic, both among habitats and within habitats, and patterns of GLS plasticity varied greatly among genotypes (Wagner and Mitchell-Olds, 2018).

*Plantago major* is an annual or facultative perennial rosette-forming herbaceous plant. Not being very competitive, *P. major* generally grows in ruderal areas especially along paths or roadsides and near gateways, where grass is short or absent (Warwick and Briggs, 1980). Native to Eurasia, *P. major* is a cosmopolitan species. It reproduces both sexually (self-compatible wind pollinated) and asexually through rosettes formation. Low genetic diversity within population of *P. major* has been shown to favour ecotypic and phenotypic differentiation (Halbritter *et al.*, 2015; Van Dijk *et al.*, 1988; Warwick and Briggs, 1980). *Plantago major* can cover a very wide elevation range: from the sea level to the alpine ecosystems all the way up to 3’000 meters above sea level (Ren *et al.*, 1999). *Plantago major* produces important amounts of secondary metabolites belonging to the class of cyclopentanoid monoterpenes called iridoids glycosides (IGs) and caffeoyl phenylethanoid glycoside (CPG) compounds (Pankoke *et al.*, 2013), which act as herbivore deterrents against generalist chewing insect (Fuchs and Bowers, 2004). IGs and CPG display a relatively high degree of variation in plant tissues depending on plant population, plant phenology and environmental factors (Barton, 2008; Bowers and Stamp, 1993; Darrow and Bowers, 1999; Darrow and Deane Bowers, 1997; Miehe-Steier *et al.*, 2015; Pellissier *et al.*, 2014), and they have been shown to display phenotypic plasticity (Bowers and Stamp, 1992; Halbritter *et al.*, 2015; Kuiper and Smid, 1985; Lotz and Blom, 1986).

### Experimental design

*Cardamine pratensis* seeds were collected from four different natural populations: two low elevation and two high elevation populations along elevation gradients of Jura Mountains in Switzerland in 2016. *Plantago major* seeds where collected from six different natural populations along three elevation gradients in the Swiss Alps during summer 2016 (Supplementary Table S1). Seeds were collected on randomly selected plants (*C. pratensis*, n= 6 plants /population; *P. major*, n= 10 plants / population) within a 100 m radius for each population. We here did not track the maternal genetic background as is classically done in selection experiment studies, because we were principally interests at ecotypic variation and not at genotypic variation. Therefore seeds within one population were pooled to obtain elevation-specific ecotypes. Seeds were germinated in Petri dishes lined with humid filter paper. One week after germination, 25 seedlings of *C. pratensis* per population (total of 100 plants) and 24 seedlings of *P. major* per population (total of 144 plants) were transplanted independently into plastic potting pots (13 cm width × 10 cm height) filled with 500 ml of sieved soil (1 cm mesh size) mixed with sand in a 3:1 ratio. Plants were immediately transferred to a climate-controlled chamber and kept at 16h/22°C - 8h/16°C day-night, and 50% relative humidity conditions for two weeks. Plants received nutrients twice a week until the beginning of reciprocal transplant experiment.

After two weeks of growth in the climate chamber, 25 *C. pratensis* plants per population and 24 *P. major* plants per population were equally distributed in two common gardens placed along the same mountain slope: La Neuveville (N: 47°06’84.28”, E: 7°10’43.9”, elevation: 450 m), and Chasseral (N: 47°07’03.36”, E: 7°01’45”, elevation: 1600 m). The plants were left growing for a period of two months during summer 2017.

### Plant growth-related traits

For both plant species, the aboveground plant parts were separated from roots at the end of the experiment, oven-dried at 40°C for 48h and weighted to determine their dry biomass. Furthermore, in *P. major* plants, two additional growth-related traits were measured. The chlorophyll content of the plant was measured as the average of three fully expanded leaves per plant using a SPAD-502Plus chlorophyll meter (Konica Minolta (China) Investment Ltd). Specific leaf area (SLA) was measured as the area (calculated using ImageJ software) of one fully expanded leaf per plant divided by their oven-dried (40°C for 48h) biomass (mm^2^ mg^−1^ DW). Higher SLA levels and chlorophyll content tend to positively correlate with potential relative growth rate across species, photosynthetic rate, or leaf nitrogen (N) (Garnier and Laurent, 1994; Poorter and Garnier, 2007). In general, species in resource-rich environments tend, on average, to have a higher SLA than do those in resource-poor environments (Garnier and Laurent, 1994; Poorter and Garnier, 2007).

### Chemical defences

All leaves were harvested immediately at the end of the field experiment prior to removal of plants from the field sites, while leaf preparation for each species followed two different methods due to the different secondary metabolite extractions and analyses.

*Cardamine pratensis* leaves were immediately frozen in liquid nitrogen and stored at − 80 °C; ground to powder using mortars and pestles in liquid nitrogen, and a 100 mg aliquot was weighed for GLS extraction. The extraction solvent (1.0 ml methanol: H_2_O: formic acid (70:29.5:0.5, v/v)) was added to the tubes along with 5 glass beads, shaken in a tissue lyser (Retsch GMBH, Haan, Germany) for 4 min at 30 Hz, and centrifuged at 12800 rpm for 3 min. The supernatant was diluted 20 times with 70% methanol and transferred to an HPLC vial. GLS identification and quantification was performed using an Acquity ultra-high pressure liquid chromatography (UHPLC) from Waters (Milford, MA) interfaced to a Synapt G2 quadrupole time-of-flight mass spectrometry (QTOF) from Waters with electrospray ionization, using the method as described in (Glauser *et al.*, 2012).

*Plantago major* leaves were oven-dried at 40 °C for 48 h prior being ground to powder using stainless steel beads in the tissue lyser, a 10 mg aliquot was weighed and a 1.5 ml methanol were added to the tubes along with 5 glass beads. The tubes were shaken 4 min at 30 Hz and centrifuged at 14000 rpm for 3 min. The supernatants were diluted five times by adding 800 µl of MilliQ water to 200 μl of pure extract. IGs and CPG were separated by UHPLC-QTOF using an Acquity BEH C18 column from Waters (50x2.1mm, 1.7 μm particle size) at a flow rate of 0.4 ml/min. The following gradient of water + formic acid 0.05% (phase A) and acetonitrile + formic acid 0.05% (phase B) was applied: 2-9 % B in 1.5 min, 9-50 % B in 3.5 min, 50-100% B in 1.5 min, held at 100% B for 1.5 min, back to 2% B and held for 2.0 min. The column was maintained at 25 °C. The injection volume was 1 μl. Detection was achieved in negative electrospray using the deprotonated ions or the formate adducts as quantification ions. Quantification ions and retention time of the two standards were: aucubin m/z 391.124 (formate adduct), retention time 1.17 min, and verbascoside m/z 623.198 (deprotonated ion), retention time 3.16 min. Absolute amounts of IGs and CPG were determined by external calibration using five standard solutions of aucubin at 0.2, 0.5, 2, 5 and 10 μg/land verbascoside at 0.2, 0.5, 2, 5 and 20 μg/ml. Concentrations were normalized to plant weight and expressed as μg/mg. Other IGs and CPG were putatively identified based on their retention time and chemical formula by comparing them to previous detection in *P. major* or in species of *Plantago* genus (Rønsted *et al.*, 2000) and database (Dictionary of Natural Products, CRC Press, USA, version 6.1. on DVD) containing information on known IGs and CPGs and quantified as aucubin or verbascoside equivalents. IGs named with the code IGs followed by numbers represent molecular formula corresponding to potential IGs for which several isomers exist in the literature and thus cannot be unequivocally annotated.

### Herbivore bioassay

To measure plant resistance against insect herbivores (resistance is defined as the effect of plant defence traits on herbivore performance (Karban and Baldwin, 1997)); we used a generalist herbivore, *Spodoptera littoralis* (Lepidoptera: Noctuidae; obtained from Syngenta, Stein AG, Switzerland). *Spodoptera littoralis* is known to feed on species belonging to more than 80 families of plants (Brown and Dewhurst, 1975), and is widely used for performing plant resistance bioassays. Newly hatched larvae were reared on corn-based artificial diet for 7 days before the beginning of the bioassay. Immediately after removal of plants from the field, both plant species were placed in a climate-controlled chamber (24 / 18 °C, 16/8 hr, day/night regime, and 55 % R.h.) to homogenize the condition for herbivores feeding on both species during bioassay performance. For *C. pratensis,* one fully expanded new leaf from 12 plant per ecotype and per population that were growing at the two elevation common gardens (n = 48) was cut and separately placed in a Petri dish on a filter paper moisten with one drop of distilled water. One 7-days old *S. littoralis* larva was added to each petri dish. For *P. major* instead, we performed a whole plant bioassay. We placed two 7-day old *S. littoralis* larvae on 24 plant per ecotype population that were growing at the two elevation common gardens (n = 96). Plants were covered with nylon nets to avoid escaping of caterpillars. After five days of herbivory for *C. pratensis* and three days for *P. major*, the insects were retrieved from individual Petri dishes and plants, respectively and their weights were measured and recorded. We consider the larval gain weight using the formula ln(*final weight - initial weight*). For *P. major* the larval gain weight represent the average of the two caterpillar placed on each plant. Lower weight gains indicate that plants are more resistant (Humphrey *et al.*, 2018).

### Statistical Analyses

All statistical analyses were performed within the R environment (R Development Core Team, 2017).

For chemical data, we calculated the sum of glucosinolate compounds (GLS total) for *C. pratensis* and the sum of iridoids glycosides (IGs total) and caffeoyl phenylethanoid glycoside (CPG total) for *P. major*, as well as a measure of chemical diversity for both plant species using the Shannon-Weaver diversity indices (Hill, 1973) with *diversity* function in the *vegan* package in R (Oksanen *et al.*, 2017).

To measure the interactive effect of transplant site and elevation of origin of the plant ecotypes on plant growth and defence traits, we used two-way ANOVAs by including transplant sites (high and low), ecotypes (high and low) and their interaction as fixed factors. We also included the term population nested within ecotypes in the model to assess variability across populations within a given elevation of origin. The response variables were; AG biomass, larval weight gain, total GLS, total indole, total aliphatic, and chemical diversity for *C. pratensis*, and AG biomass, chlorophyll content, SLA, larval weight gain, total chemistry, total IGs, total CPG and chemical diversity for P. *major*. All chemical traits were log-transformed prior analyses to meet normality and homoscedasticity assumptions. A significant effect of site of growth (i.e. elevation) would indicate a plastic response to different environmental conditions. A significant effect of ecotype would indicate differentiation in traits among populations belonging to different ecotypes. A significant effect of population would indicate differentiation in traits among populations. A significant elevation × ecotype term would indicate ecotype-specific selection for plasticity for a given trait.

To address the multivariate nature of plant secondary compounds, we also ran a full-factorial model including the individual secondary metabolites abundance matrix as response variable and plant ecotype and elevation as factors using permutational analysis of variance (PERMANOVA) with the *adonis* function in the package vegan in R (Oksanen *et al.*, 2017). We also included plant biomass as covariate to control for potential direct effect of biomass on plant chemistry (Züst *et al.*, 2015). The Bray–Curtis metric was used to calculate a dissimilarity matrix of all compounds among samples for the PERMANOVA. We visualized ecotypic differentiation of the secondary metabolites using an NMDS ordination analysis of the chemical compounds based on Bray Curtis distance (package vegan in R) (Oksanen *et al.*, 2017).

Finally, to visualize and calculate the magnitude of plasticity of the plant growth and defence related traits when plants were places in their non-native habitat, we calculated effect sizes for all traits as the log-response-ratio 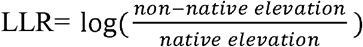 using the *effsize* function in the *effsize* package in R (Torchiano, 2017), and when significant, we reported them as standardized mean difference (SMD) values. The figure constructed based on effect size aims at representing the plastic response of traits, G×E effects, as well as the magnitude of responses. A 95% of confidence interval bar that deviates from zero shows a significant effect of treatment (positive or negative effect of non-native growing elevation) (Nakagawa and Cuthill, 2007), while a deviation of one of the interval bars from zero, but not the other, indicates G×E effects.

## Results

### Plant growth related traits

For both species, we observed phenotypic plasticity and ecotypic differentiation in aboveground (AG) biomass through significant effects of both ecotype and elevation (high or low elevation growing sites) (Fig. 2, 3, 4; Table 1). We observed that AG biomass of high elevation ecotypes increased by 49% (SMD = 1.17) for *C. pratensis* and by 45% (SMD = 1.48) for *P. major* at the non-native elevation (low elevation site), while low elevation ecotypes’ AG biomass decreased by 61% (SMD = - 0.96) for *C. pratensis* and by 51% (SMD = - 1.93) for *P. major* at the non-native elevation (high elevation site) (Fig. 2, 3, 4; Table 1). Furthermore, our results indicated that high elevation ecotypes produced 38.5 % and 12% more AG biomass than low elevation ecotypes in *C. pratensis* and *P. major*, respectively. In addition, in *P. major* leaf chlorophyll content and SLA showed plasticity through growing elevation effect, with the latter also showing marginal G×E effect. Specifically, we observed that chlorophyll content of high elevation ecotypes increased by 4.1% (SMD = 1.55) at the non-native site (low elevation site) and low elevation ecotypes had 3.4% (SMD = −1.36) less chlorophyll content at the non-native site (high elevation) (Fig. 2B, 4; Table 1). Moreover, SLA of low elevation ecotypes significantly increased by 6.6% (SMD = 0.96) at their non-native growing site (Fig. 2B, 4; Table 1).

**Fig. 2.**
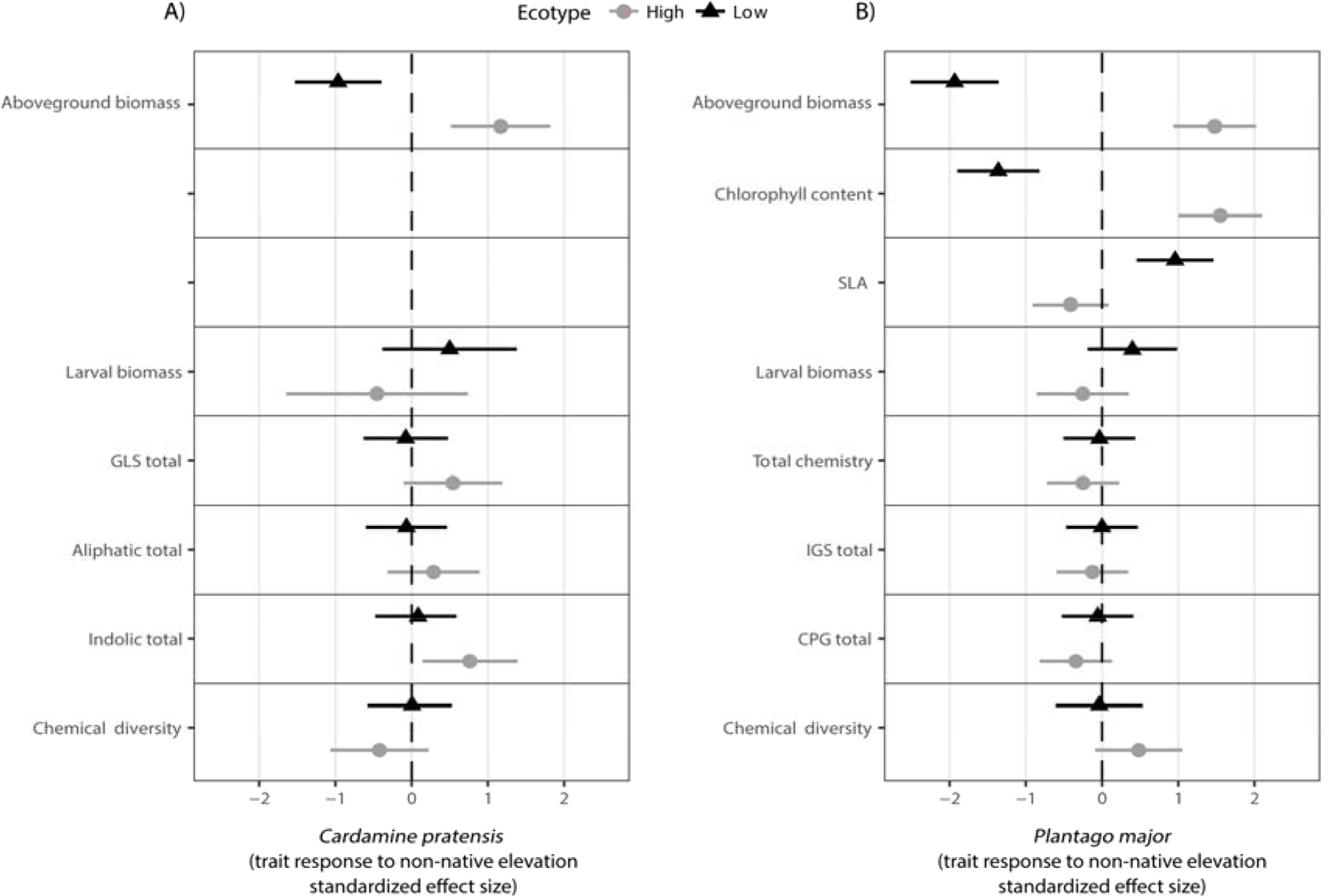
Effect sizes for the influence of non-native growing elevation on plant growth and defence related trait for high and low elevation ecotypes of *C pratensis* (A) and *P. major* (B). Effects are natural log response ratios (LRRs) with 95% confidence limits.

**Table 1.**
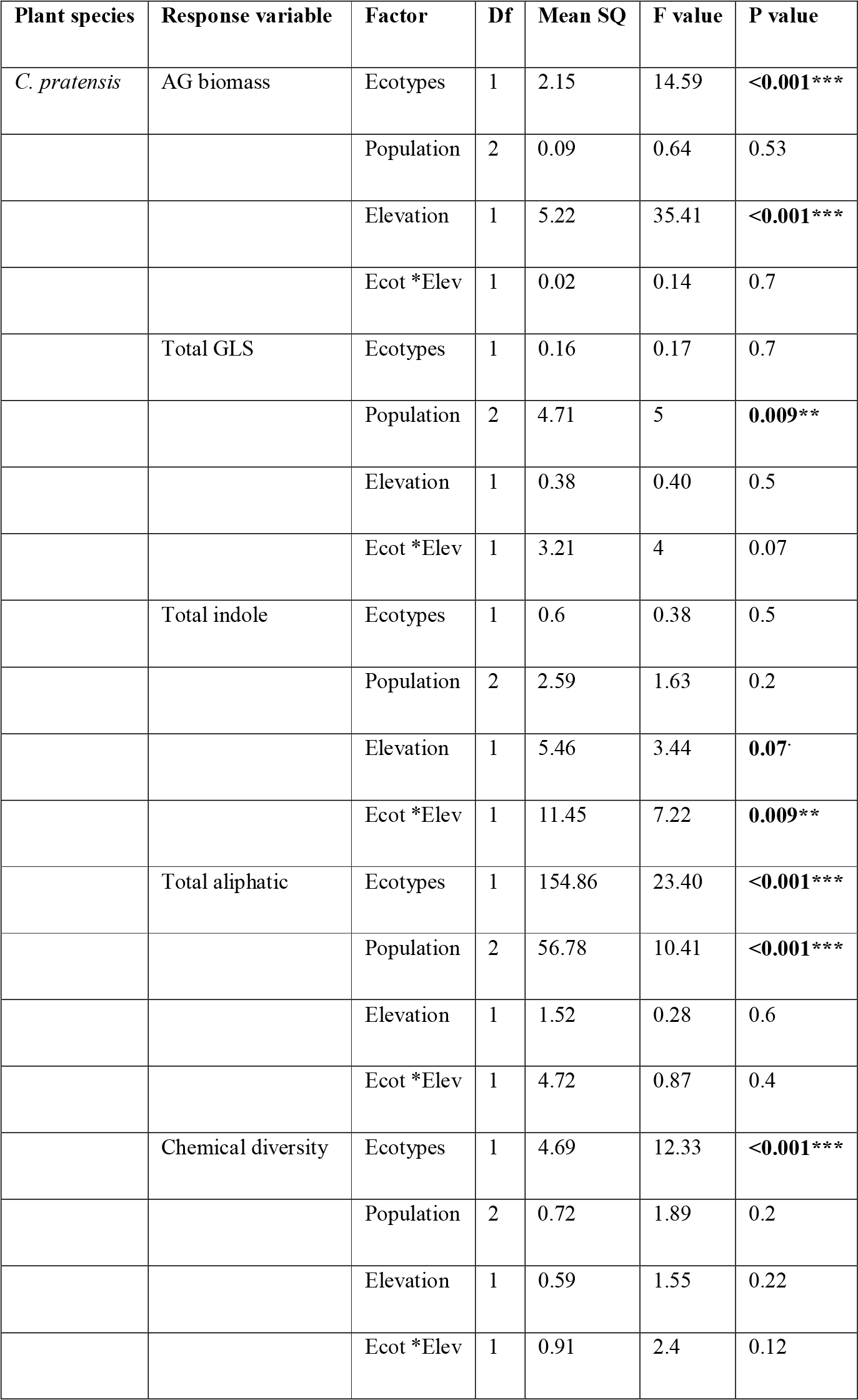
Two-way ANOVA Table for measuring the interaction between the effects of high and low elevation ecotypes and the elevation of growth in two common garden sites on growth and defence traits.

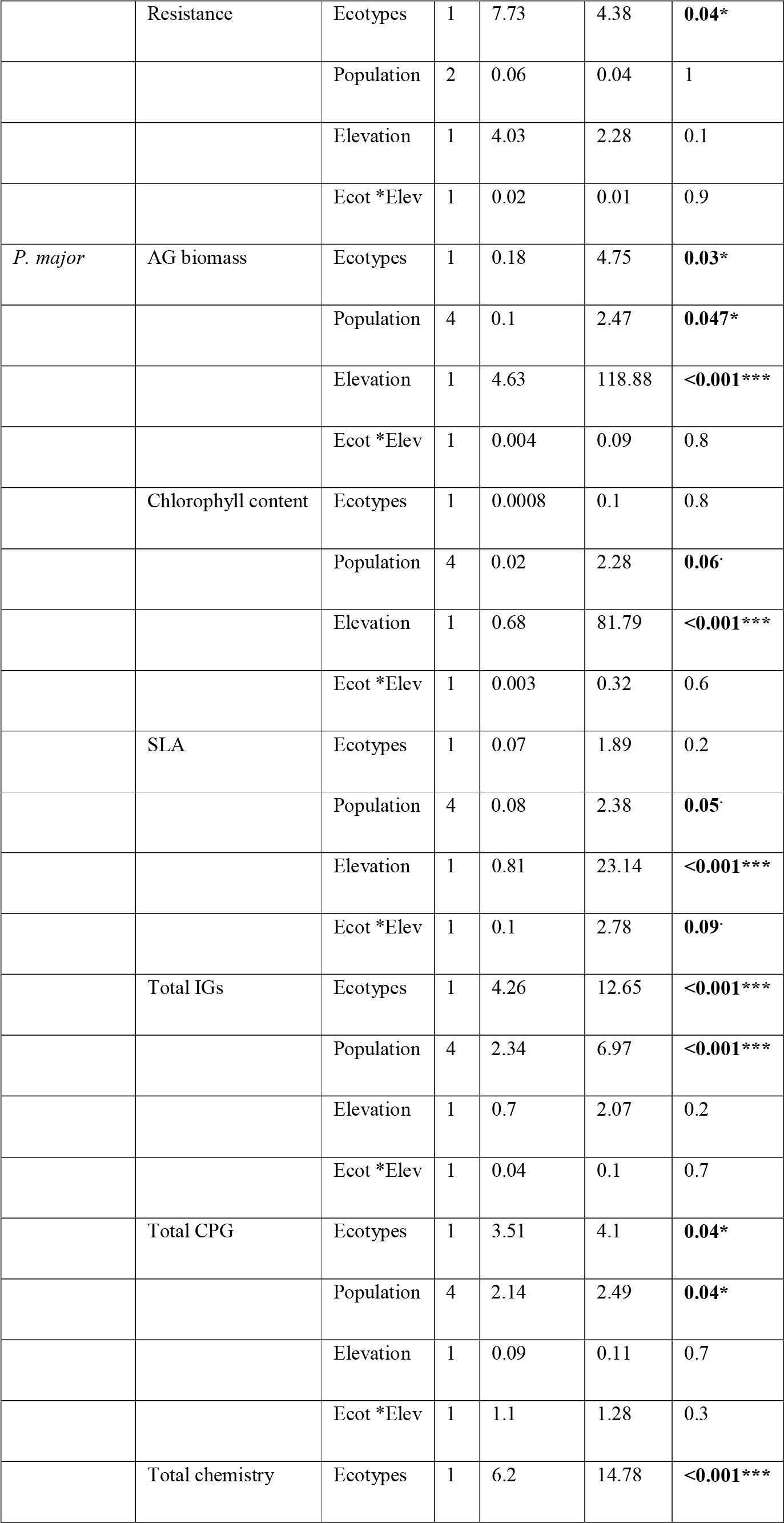

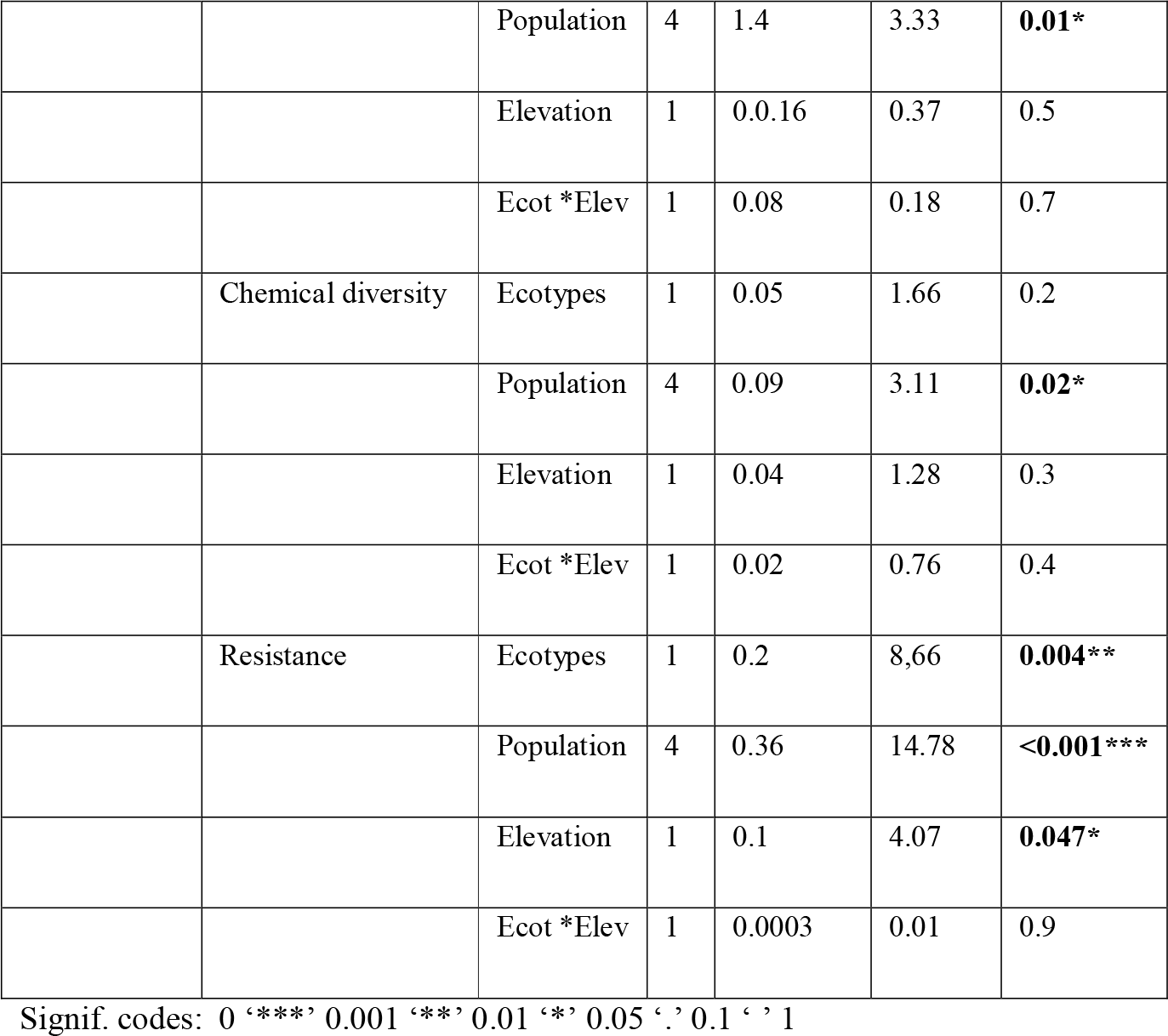

### Plant chemical defences and resistance

The GLS profiles of *C. pratensis* leaves consisted of six GLS compounds (two aliphatic, three indoles and one aromatic), and the secondary metabolites profile of the *P. major* leaves consisted of 13 IGs and 3 CPG compounds (Supplementary Fig. S2). In *C. pratensis*, we observed phenotypic plasticity in total indole GLS, specifically through significant ecotype by elevation interaction (G×E effect), where the total indole GLS concentration in high elevation ecotypes significantly increased at the low elevation site (non-native) by 28% (SMD = 0.77) (Fig. 2A, 3; Table 1). Moreover, we found ecotypic effect for *S. littoralis* larval weight gain; larvae on low elevation ecotypes grew 81% more compared to high elevation ecotypes. Low elevation ecotypes produced 37% more aliphatic GLS than high elevation ecotypes, and high elevation ecotypes showed 25% more chemical diversity than low elevation ecotypes (Fig. 3, Table 1). Furthermore, the PERMANOVA showed that the abundance and chemical diversity of GLS were globally affected by plant ecotypes (P= 0.001, Fig. 5A-B). In *P. major,* we also found ecotypic differentiation for *S. littoralis* larval weight gain; larvae on low elevation ecotypes grew 8% more than on high elevation ecotypes. Low elevation ecotypes produced 17%, 17% and 22% more total chemistry; total IGs and total CPG than high elevation ecotypes, respectively (Fig. 4, Table 1). The PERMANOVA revealed plant ecotypic effect (P= 0.001) and growing elevation effect (P= 0.005) (Fig. 5C-D) in the abundance and diversity of secondary metabolites in *P. major*. Additionally, we found that abundance of the total chemistry and diversity of the compounds were significantly affected by the AG biomass of *P. major* (P= 0.0002).

**Fig. 3.**
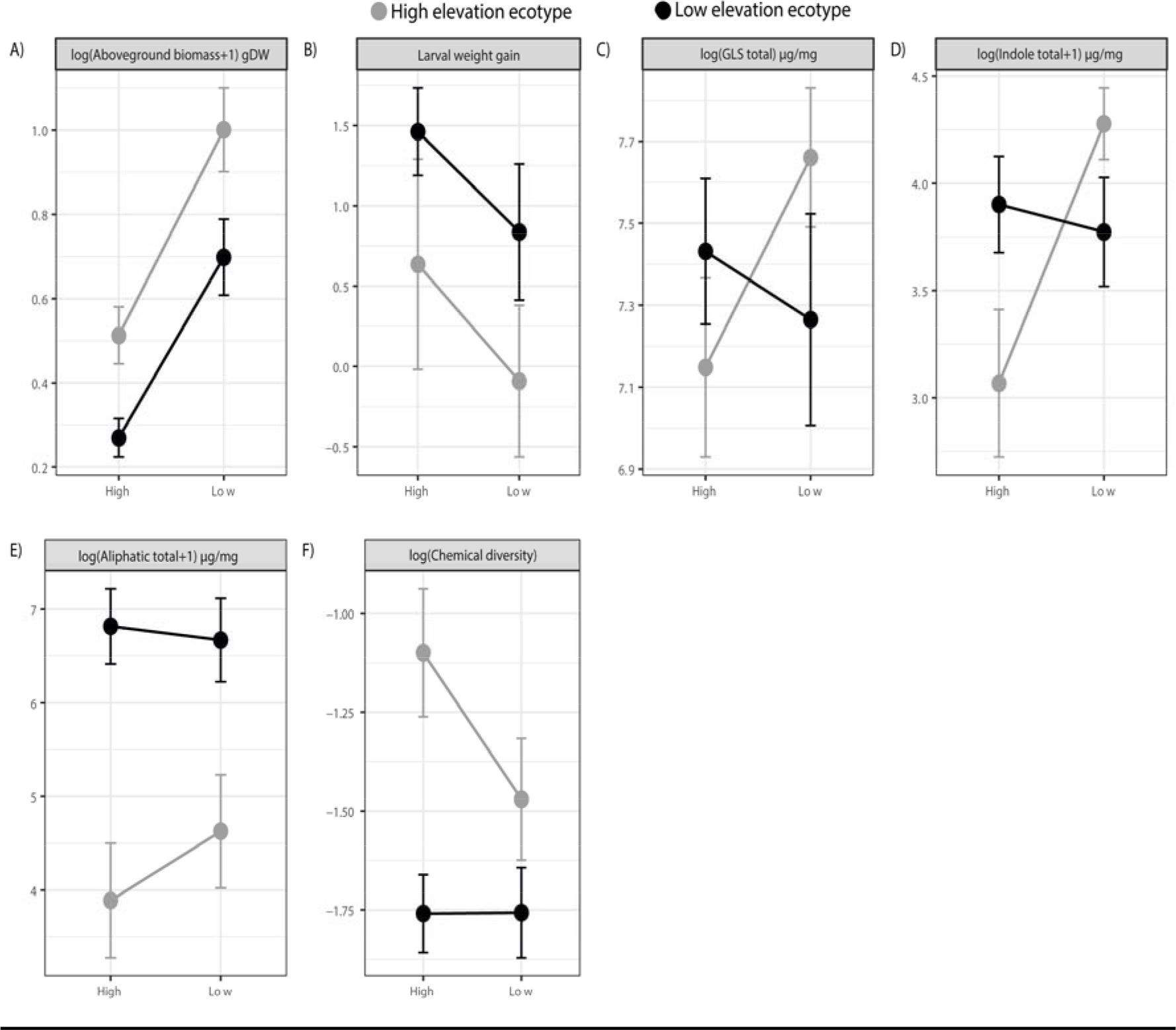
Reaction norms of *C. pratensis* ecotypes of growth (A), resistance (B) and defence (C, D, E, F) traits. Mean phenotypic values (mean ± 1 s.e. for each elevation ecotype) are represented in black (low elevation ecotypes) and in grey (high elevation ecotypes) across two contrasted growing elevations (high or low elevation).

**Fig. 4.**
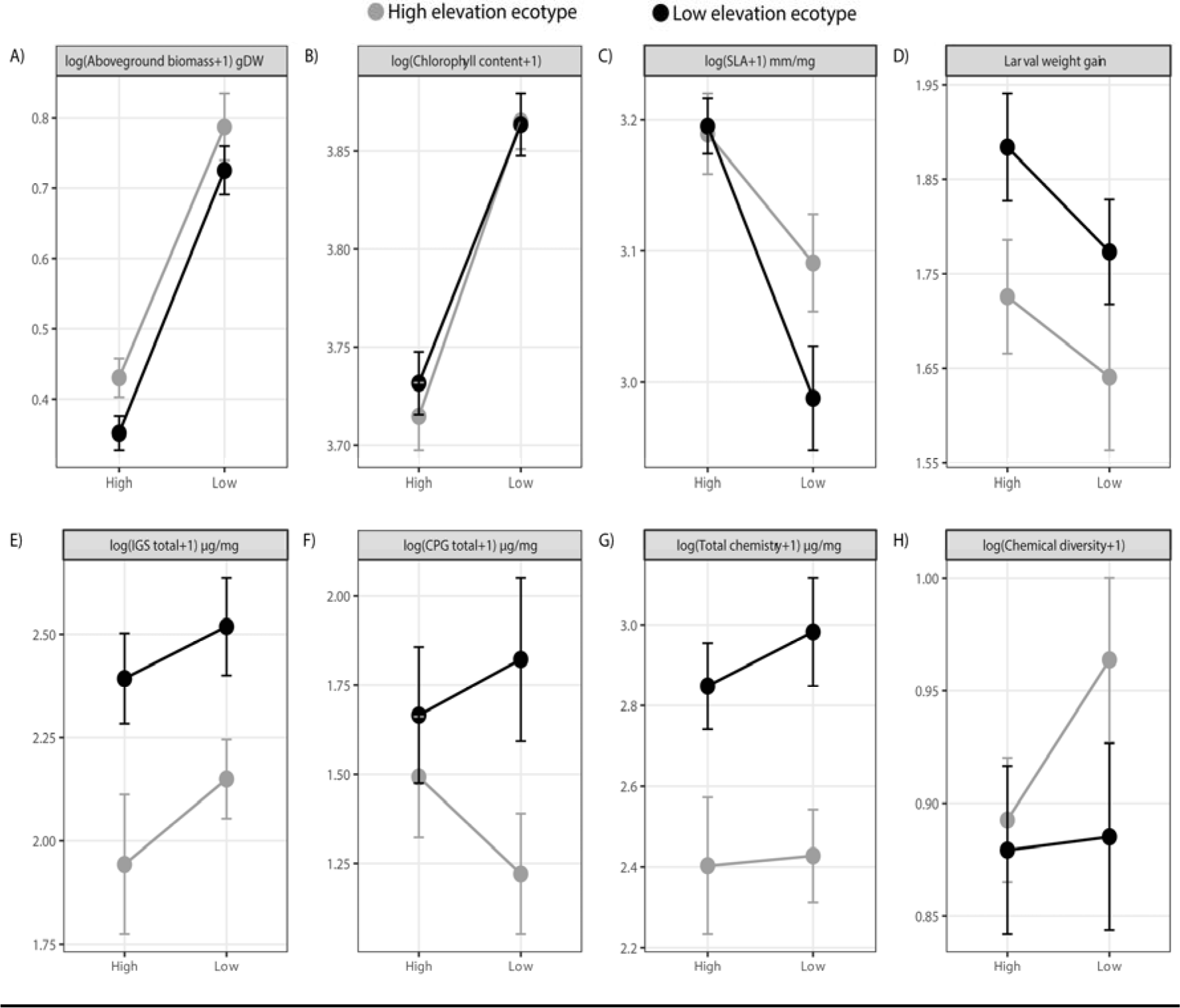
Reaction norms of *P. major* ecotypes of growth (A, B, C), resistance (D) and defence (E, F, G, H) traits. Mean phenotypic values (mean ± 1 s.e. for each elevation ecotype) are represented in black (low elevation ecotypes) and in grey (high elevation ecotypes) across two contrasted growing elevations (high or low elevation).

**Fig. 5.**
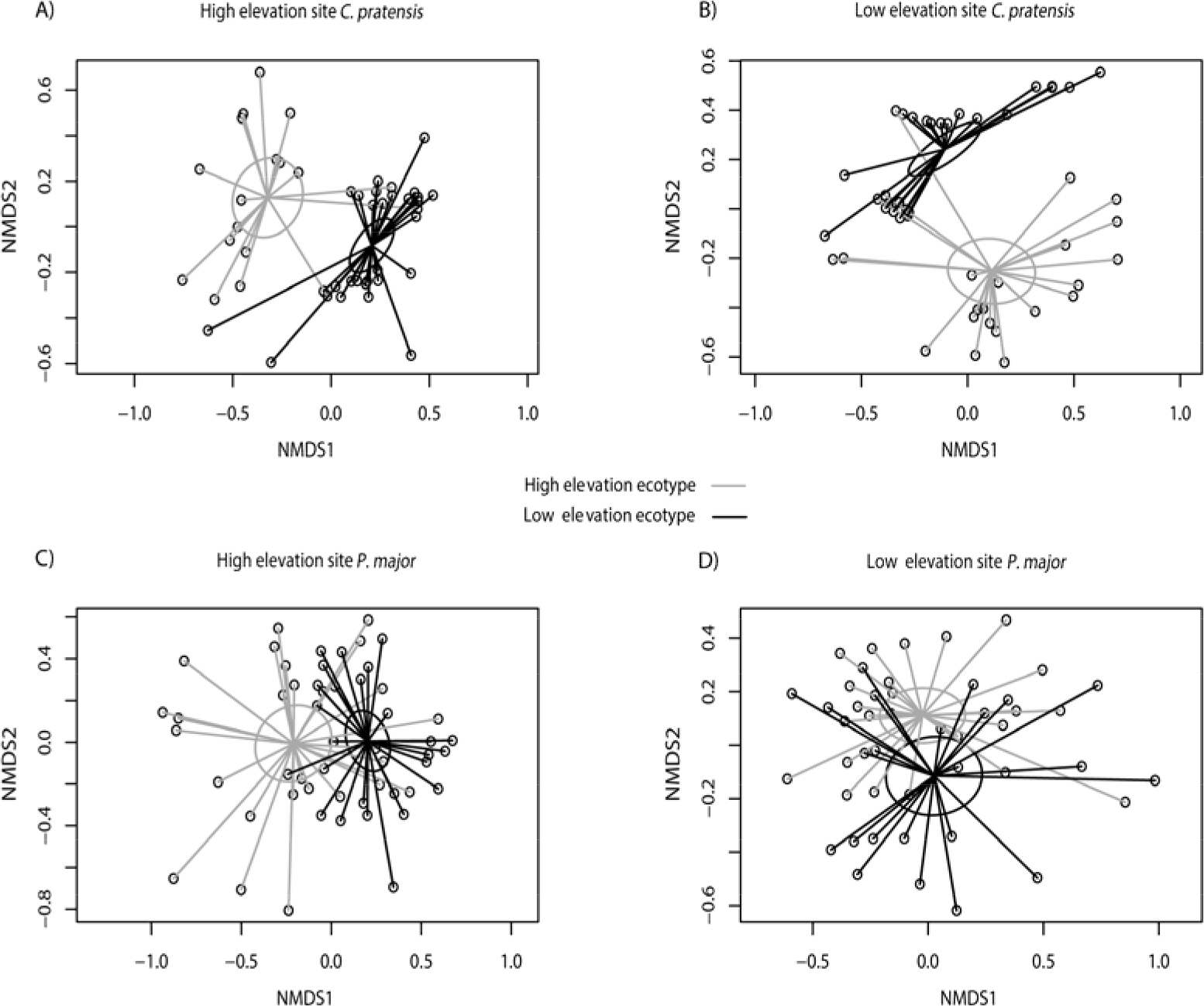
Glucosinolates (GLS), *iridoid glycosides* (*IGs*) *and* caffeoyl phenylethanoid glycoside (CPG) ordinations. Representation of the non-multidimensional scaling (NMDS) indicating the GLS found in high and low *C. pratensis* ecotypes at high (A) and low (B) elevation sites and IGs and CPG found in *P. major* ecotypes at high (C) and low (D) elevation sites. The 95% confidence interval ellipses are represented based on the two elevation ecotypes (high elevation ecotype in grey and low elevation ecotype in black). Stress values: (A) and (B) = 0.12, (C) and (D) = 0.2, K = 2.

Overall, we also found significant population-level effects in trait expression. For instance, we found a significant effect of plant population for *C. pratensis* total GLS and aliphatic GLS (Supplementary Fig. S3 and Table 1). In *P. major,* we observed significant effects of plant population on all the measured traits (marginal for SLA and chlorophyll content) (Supplementary Fig. S4 and Table 1).

## Discussion

Using reciprocal transplant experiments of ecotypes growing at different elevation, we observed ecotypic differentiation accompanied by plasticity in growth related traits, while we mainly observed ecotypic differentiation for defence and resistance traits for both *P. major* and *C. pratensis*. Below, we outline the potential causes for such divergence along elevation gradients.

### Plant biomass accumulation

We found high levels of phenotypic plasticity in the observed AG production pattern. Plasticity can be visualized as a change in the slope of the reaction norm between the ancestral and derived population or species (Doughty, 1995; Gotthard *et al.*, 1995). In this regard, for both species plant growth related traits (plant biomass, leaf chlorophyll content and SLA) showed plasticity. Our results compliment other findings where the combination of ecotypic differentiation and phenotypic plasticity in growth-related traits such as biomass and flower size was shown for invasive species at their invasive range (Martín-Forés *et al.*, 2017). More specifically, we observed that in both species, the AG biomass across both ecotypes increased at low elevation growing site and decreased at high elevation growing site. Increase in AG biomass of both ecotypes at low elevation growing site comes as no surprise, given the growing condition at low elevation are warmer and more favourable than at high elevation. Two reasons have been put forward for plants to reduce growth at high elevation. First, a decrease in the general metabolic activity as a function of colder temperature inhibits photosynthetic rate and biomass production (Boyer, 1982). Second, it has been proposed that because plants growing at higher elevations typically receive direct sunlight and higher ultraviolet radiation, and ultraviolet radiation destroys the auxins content at the apical shoots, they tend to grow much slower than lowland plants (Keller *et al.*, 2004). Furthermore, both *C. pratensis* and *P. major* are perennial species and it can be argued that high elevation ecotypes accumulated higher AG biomass than low elevation ecotypes once placed in more favourable conditions of low elevation to compensate for the next year’s growing season when they would have to allocate more resource to flower and seed production. Such a scenario should be less likely for low elevation plants growing at their native site.

Interestingly, we also observed that high elevation ecotypes produced more biomass than low elevation ecotypes, and this was true for both species. This is somewhat surprising since we expected alpine plants to grow smaller in harsher and colder environments (Atkin and Day, 1990; Körner, 2003). While plant size is negatively correlated to extremely cold temperatures (Squeo *et al.*, 1991) and, as a consequence, generally decreases with elevation (Körner, 2003), it appears that high-elevation ecotypes favour fast biomass accumulation (Körner, 2016).

Plants adapted to growing in cold conditions, such as in high altitude climates, where growing season is short, pass through seasonal development taking advantage of the warmest period of the growing season. In addition, plants growing at cold condition typically exhibit greater photosynthetic and respiratory capacities than their warm-grown counterparts (Atkin *et al.*, 2006).Therefore, high-elevation ecotypes could highly benefit from faster development and high rates of metabolism (Körner, 2016), and, at equal growing conditions (same soil) and during the same growing period, can accumulate more biomass than their low-elevation counterparts.

### Plant chemical defences and resistance

Concerning plant defences and resistance, we observed ecotypic differentiation across most defence and resistance measures, including total GLS, aliphatic GLS, chemical diversity, total IGs, total CPG, total chemistry and larval weight. Generally, regardless of the growing elevation, low-elevation ecotypes produced more chemical defences. The strong effect of temperature on plant primary and secondary metabolism is known and our results are in line with other findings showing the temperature-driven suppression of plant secondary metabolites at high elevation (Pellissier *et al.*, 2014) and general decrease in secondary metabolite production from low to high elevation (Kergunteuil *et al.*, 2018). However, a decrease in secondary metabolite production at high elevation could also be attributed to a relaxation of herbivory pressure at high elevation (Pellissier *et al.*, 2014). To date we have no data that allows disentangling biotic and abiotic effects of defence decline at high elevation. Interestingly, however, indole GLS showed no ecotypic differentiation, in which, high elevation ecotypes produced more of these compounds when placed at low elevation (see G x E effect in Table 1).

Unlike aliphatic GLS, for which induction has been rarely observed (Koritsas *et al.*, 1991; Li *et al.*, 1999), induction of indolic GLS has been wildly documented in several systems (Agrawal *et al.*, 1999; Doughty *et al.*, 1995; Griffiths *et al.*, 1994; Moyes *et al.*, 2000; Raybould and Moyes, 2001; Siemens and Mitchell-Olds, 1998), including the closely related *Cardamine hirsuta* (Bakhtiari et al., unpublished data). Additionally, in contrast to the aliphatic GLS that are under strong genetic control (Raybould and Moyes, 2001), indole GLS have been shown to be strongly influenced by environmental factors with some heritable variation in production (Rücker and Röbbelen, 1994). Altogether, this could indicate that the production of indole GLS might be more costly than the production of other GLS (Bidart-Bouzat *et al.*, 2005; Traw, 2002) in *C. pratensis.* Several studies detected a relatively high cost of plasticity for chemical defences and emphasized on the fact that the limit of plasticity in expression of chemical traits may be attributed to such cost (Agrawal *et al.*, 2002; Agrawal *et al.*, 2010; Züst and Agrawal, 2017). On the other hand, plasticity in defence-related traits is the reflection of both biotic and abiotic environmental conditions that affect the expression of defences, and plasticity of defence-related traits in response to biotic pressures, such as herbivory, is well-documented (Agrawal *et al.*, 2002; Humphrey *et al.*, 2018; Wagner and Mitchell-Olds, 2018). Such results may suggest that plants show higher degree of plastic response to biotic stimuli compared to abiotic stress such as environmental fluctuations. Thus, the lack of plasticity in the majority of the measured defence-related traits in our study could be due to the fact that the benefits of plasticity in expression of defence cannot outweigh the costs of biotic pressure occurred early in the season or other potential costs of defence plasticity. For example, indolic GLS were not plastic, in contrast to plastic non-indolic GLS, in *Cardamine cordifolia* plants growing in shaded-common gardens, that are characterized by low herbivory (Humphrey *et al.*, 2018). In contrast to our results, in the same study, Humphrey et al. found plasticity in larval weight gain of specialist herbivore (*Scaptomyza nigrita*).

On a broader perspective, detailed analysis of the effect sizes (ESs) between growth and defence related traits in *C. pratensis* indicates that the magnitude of plastic responses displayed by high elevation ecotypes is higher for AG biomass (very large ES) compared to indolic GLS production (large ES). In *P. major* the magnitude of plastic responses in all the growth-related traits were also very large, compared to the non-significant plastic responses for all the defence-related traits (except for some the individual compounds, Supplementary Fig. S2B). Nevertheless, the lack of plastic response to elevation in defence-related traits does not rule out the potential for plasticity in chemical defences. Given the fact that the environmental effect of the growing elevation could affect the plant chemistry at any time throughout the growing season and the chemistry was measured only at the end of the field season, a potential plasticity in expression of such traits could have disappeared by the end of the season. Moreover, detecting plastic response in defence traits upon biotic stress such as herbivory is simpler. Upon herbivory, the phytohormone activation machinery behind expression of chemical defences is an immediate process, whereas the detection of the potential plastic responsiveness of plant defence to abiotic stimuli might be masked by the time-dependency of the growing season. For instance, phenotypic plasticity in flowering time in response to seasonal variations has been shown both in controlled environment (Anderson *et al.*, 2011) as well as from long-term field survey in *Boechera stricta* (Anderson *et al.*, 2012). In addition, two studies, one on *C. cordifolia* and the second on *P. lanceolata*, showed phenological variation in GLS and IGs plant tissue content, respectively (Darrow and Deane Bowers, 1997; Rodman and Louda, 1984). Therefore, ontogeny should also be addressed when measuring plasticity because plants have been shown to express different levels of plasticity in defence traits as they grow.

## Conclusions

Few studies have assessed phenotypic variation of plant growth versus defence traits in response to contrasting environments. Here, we document that plant growth traits displayed strong ecotypic differentiation accompanied by plasticity, but, in contrast, we find little support of phenotypically plastic defence and resistance traits in response to different growing habitat across step elevation gradient. Future research on similar systems would require coupling the observed effects on plant phenotypes with fitness measurements and selection gradient analyses in order to disentangle the fitness benefits of phenotypic plasticity versus fixed ecotypic differentiation for individual plant traits.

## Supplementary Data

**Table S1** Coordinates of the populations of *C. pratensis* and *P. major*.

**Fig. S1.** Natural distribution map of *C. pratensis* and *P. major* along elevation.

**Fig. S2.** Effect sizes of individual chemical compounds of *C. pratensis* and *P. major* growing at their non-native elevation.

**Fig. S3.** Reaction norms of growth and defense traits for populations of *C. pratensis*.

**Fig. S4.** Reaction norms of growth and defense traits for populations of *P. major*.

## Acknowledgments

We thank Adrienne Godschalx for her valuable comments on the manuscript. This work was financed by a Swiss National Science Foundation grant 159869 to SR.

## References

Aeschimann D, Lauber K, Moser DM, Theurillat J-P. 2004. Flora alpina: ein Atlas sämtlicher 4500 Gefässpflanzen der Alpen.

Agrawal AA, Conner JK, Johnson MTJ, Wallsgrove R. 2002. Ecological genetics of an induced plant defense against herbivores: Additive genetic variance and costs of phenotypic plasticity. Evolution 56, 2206–2213.

Agrawal AA, Conner JK, Rasmann S. 2010. Tradeoffs and negative correlations in evolutionary ecology. Evolution since Darwin; the first 150 years, Vol. 150. Stony Brook, NY: Sinauer Associates, 243–268.

Agrawal AA, Strauss SY, Stout MJ. 1999. Costs of induced responses and tolerance to herbivory in male and female fitness components of wild radish. Evolution 53, 1093–1104.

Anderson JT, Inouye DW, McKinney AM, Colautti RI, Mitchell-Olds T. 2012. Phenotypic plasticity and adaptive evolution contribute to advancing flowering phenology in response to climate change. Proceedings of the Royal Society B: Biological Sciences 279, 3843–3852.

Anderson JT, Lee C-R, Mitchell-Olds T. 2011. Life history QTLs and natural selection on flowering time in Boechera stricta, a perennial relative of Arabidopsis. Evolution; international journal of organic evolution 65, 771–787.

Atkin O, Day D. 1990. A comparison of the respiratory processes and growth rate of selected australian alpine and related lowland plant species. Functional Plant Biology 17, 517–526.

Atkin OK, Loveys BR, Atkinson LJ, Pons TL. 2006. Phenotypic plasticity and growth temperature: understanding interspecific variability. Journal of Experimental Botany 57, 267–281.

Baker HG. 1974. The evolution of weeds. Annual Review of Ecology and Systematics 5, 1-24. Baldwin JM. 1896. A new factor in evolution. The American Naturalist 30, 441–451.

Baldwin JM. 1896. A new factor in evolution. The American Naturalist 30, 441–451.

Barton KE. 2008. Phenotypic plasticity in seedling defense strategies: compensatory growth and chemical induction. Oikos 117, 917–925.

Bidart-Bouzat MG, Mithen R, Berenbaum MR. 2005. Elevated CO2 influences herbivory-induced defense responses of Arabidopsis thaliana. Oecologia 145, 415–424.

Bolser RC, Hay ME. 1996. Are tropical plants better defended? Palatability and defenses of temperate vs. tropical seaweeds. Ecology 77, 2269–2286.

Bowers MD, Stamp NE. 1992. Chemical variation within and between individuals of Plantago lanceolata (Plantaginaceae). Journal of Chemical Ecology 18, 985–995.

Bowers MD, Stamp NE. 1993. Effects of plant age, genotype and herbivory on Plantago performance and chemistry. Ecology 74, 1778–1791.

Boyer JS. 1982. Plant productivity and environment. Science 218, 443–448.

Bradshaw AD. 1965. Evolutionary significance of phenotypic plasticity in plants. In: Caspari EW, Thoday JM, eds. Advances in Genetics, Vol. 13: Academic Press, 115–155.

Brown ES, Dewhurst CF. 1975. The genus spodoptera (Lepidoptera, Noctuidae) in Africa and the Near East. Bulletin of Entomological Research 65, 221–262.

Callis-Duehl K, Vittoz P, Defossez E, Rasmann S. 2017. Community-level relaxation of plant defenses against herbivores at high elevation. Plant Ecology 218, 291–304.

Chapin FS, Chapin MC. 1981. Ecotypic differentiation of growth processes in Carex aquatilis along latitudinal and local gradients. Ecology 62, 1000–1009.

Coley PD, Aide TM. 1991. Comparison of herbivory and plant defenses in temperate and tropical broad-leaved forests. In: Price PW, Lewinsohn TM, Fernandes GW, Benson WW, eds. Plant-animal interactions: evolutionary ecology in tropical and temperate regions. New York: Wiley, 25–49.

Coley PD, Barone JA. 1996. Herbivory and plant defenses in tropical forests. Annual Review of Ecology and Systematics 27, 305–335.

Darrow K, Bowers MD. 1999. Effects of herbivore damage and nutrient level on induction of iridoid glycosides in plantago lanceolata. Journal of Chemical Ecology 25, 1427–1440.

Darrow K, Deane Bowers M. 1997. Phenological and population variation in iridoid glycosides of Plantago lanceolata (Plantaginaceae). Biochemical Systematics and Ecology 25, 1–11.

Daxenbichler ME, Spencer GF, Carlson DG, Rose GB, Brinker AM, Powell RG. 1991. Glucosinolate composition of seeds from 297 species of wild plants. Phytochemistry 30, 2623–2638.

Defossez E, Pellissier L, Rasmann S. 2018. The unfolding of plant growth form-defence syndromes along elevation gradients. Ecology Letters 21, 609–618.

Descombes P, Marchon J, Pradervand J-N, Bilat J, Guisan A, Rasmann S, Pellissier L. 2016. Community-level plant palatability increases with elevation as insect herbivore abundance declines. Journal of Ecology 105, 142–151.

Dobzhansky T. 1950. Evolution in the tropics. American Scientist 38, 209–221.

Doughty KJ, Kiddle GA, Pye BJ, Wallsgrove RM, Pickett JA. 1995. Selective induction of glucosinolates in oilseed rape leaves by methyl jasmonate. Phytochemistry 38, 347–350.

Doughty P. 1995. Testing the ecological correlates of phenotypically plastic traits within a phylogenetic framework. Acta Oecologica 16, 519–524.

Fuchs A, Bowers MD. 2004. Patterns of iridoid glycoside production and induction in plantago lanceolata and the importance of plant age. Journal of Chemical Ecology 30, 1723–1741.

Garnier E, Laurent G. 1994. Leaf anatomy, specific mass and water content in congeneric annual and perennial grass species. New Phytologist 128, 725–736.

Ghalambor CK, Mckay JK, Carroll SP, Reznick DN. 2007. Adaptive versus non-adaptive phenotypic plasticity and the potential for contemporary adaptation in new environments. Functional Ecology 21, 394–407.

Giamoustaris A, Mithen R. 1995. The effect of modifying the glucosinolate content of leaves of oilseed rape (Brassica napus ssp. oleifera) on its interaction with specialist and generalist pests. Annals of Applied Biology 126, 347–363.

Glauser G, Schweizer F, Turlings TC, Reymond P. 2012. Rapid profiling of intact glucosinolates in Arabidopsis leaves by UHPLC-QTOFMS using a charged surface hybrid column. Phytochem Anal 23, 520–528.

Gotthard K, Nylin S, xf, ren. 1995. Adaptive plasticity and plasticity as an adaptation: A selective review of plasticity in animal morphology and life history. Oikos 74, 3–17.

Gratani L, Meneghini M, Pesoli P, Crescente MF. 2003. Structural and functional plasticity of Quercus ilex seedlings of different provenances in Italy. Trees 17, 515–521.

Griffiths DW, Birch ANE, Macfarlane-Smith WH. 1994. Induced changes in the indole glucosinolate content of oilseed and forage rape (Brassica napus) plants in response to either turnip root fly (Delia floralis) larval feeding or artificial root damage. Journal of the Science of Food and Agriculture 65, 171–178.

Halbritter AH, Billeter R, Edwards PJ, Alexander JM. 2015. Local adaptation at range edges: comparing elevation and latitudinal gradients. Journal of Evolutionary Biology 28, 1849–1860.

Hill MO. 1973. Diversity and evenness: A unifying notation and its consequences. Ecology 54, 427–432.

Hopkins RJ, Ekbom B, Henkow L. 1998. Glucosinolate content and susceptibility for insect attack of three populations of Sinapis alba. Journal of Chemical Ecology 24, 1203–1216.

Hufford KM, Mazer SJ. 2003. Plant ecotypes: genetic differentiation in the age of ecological restoration. Trends in Ecology & Evolution 18, 147–155.

Hultén E, Fries M. 1986. Atlas of North European vascular plants north of the tropic of cancer: Koeltz Scientific.

Humphrey PT, Gloss AD, Frazier J, Nelson-Dittrich AC, Faries S, Whiteman NK. 2018. Heritable plant phenotypes track light and herbivory levels at fine spatial scales. Oecologia 187, 427–445.

Karban R, Baldwin IT. 1997. Induced responses to herbivory / Richard Karban and Ian T. Baldwin: Chicago : The University of Chicago Press.

Kawecki TJ, Ebert D. 2004. Conceptual issues in local adaptation. Ecology Letters 7, 1225–1241.

Keller CP, Stahlberg R, Barkawi LS, Cohen JD. 2004. Long-term inhibition by auxin of leaf blade expansion in bean and Arabidopsis. Plant Physiology 134, 1217–1226.

Kergunteuil A, Descombes P, Glauser G, Pellissier L, Rasmann S. 2018. Plant physical and chemical defence variation along elevation gradients: a functional trait-based approach. Oecologia 187, 561–571.

Kliebenstein D, Pedersen D, Barker B, Mitchell-Olds T. 2002. Comparative analysis of quantitative trait loci controlling glucosinolates, myrosinase and insect resistance in Arabidopsis thaliana. Genetics 161, 325–332.

Kliebenstein DJ, Kroymann J, Brown P, Figuth A, Pedersen D, Gershenzon J, Mitchell-Olds T. 2001. Genetic control of natural variation in Arabidopsis glucosinolate accumulation. Plant Physiology 126, 811–825.

Koritsas V, Lewis J, Fenwick G. 1991. Glucosinolate responses of oilseed rape, mustard and kale to mechanical wounding and infestation by cabbage stem flea beetle (Psylliodes chrysocephala). Annals of Applied Biology 118, 209–221.

Körner C. 2003. Alpine plant life: functional plant ecology of high mountain ecosystems. Berlin: Springer.

Körner C. 2007. The use of ‘altitude’ in ecological research. Trends in Ecology & Evolution 22, 569–574.

Körner C. 2016. Plant adaptation to cold climates. F1000Research 5, F1000 Faculty Rev-2769.

Kuiper D, Smid A. 1985. Genetic differentiation and phenotypic plasticity in Plantago major ssp major. I. The effect of differences in level of irradiance on growth, photosynthesis, respiration and chlorophyll content. Physiologia Plantarum 65, 520–528.

Lambrix V, Reichelt M, Mitchell-Olds T, Kliebenstein DJ, Gershenzon J. 2001. The Arabidopsis Epithiospecifier Protein Promotes the Hydrolysis of Glucosinolates to Nitriles and Influences Trichoplusia ni Herbivory. Plant Cell 13, 2793–2808.

Leggett HC, Brown SP, Reece SE. 2014. War and peace: social interactions in infections. Philosophical Transactions of the Royal Society B: Biological Sciences 369.

Li Y, Kiddle G, Bennett R, Wallsgrove R. 1999. Local and systemic changes in glucosinolates in Chinese and European cultivars of oilseed rape (Brassica nap us L.) after inoculation with Sclerotinia sclerotiorum (stem rot). Annals of Applied Biology 134, 45–58.

Lortie CJ, Aarssen LW. 1996. The specialization hypothesis for phenotypic plasticity in plants. International Journal of Plant Sciences 157, 484–487.

Lotz LAP, Blom CWPM. 1986. Plasticity in life-history traits of Plantago major L. ssp. pleiosperma Pilger. Oecologia 69, 25–30.

Louda SM, Rodman JE. 1983. Ecological patterns in the glucosinolate content of a native mustard,Cardamine cordifolia, in the rocky mountains. Journal of Chemical Ecology 9, 397–422.

Lövkvist B. 1956. The Cardamine pratensis complex: outlines of its cytogenetics and taxonomy, Acta Universitatis Upsaliensis.

Martín-Forés I, Avilés M, Acosta-Gallo B, Breed MF, del Pozo A, de Miguel JM, Sánchez-Jardón L, Castro I, Ovalle C, Casado MA. 2017. Ecotypic differentiation and phenotypic plasticity combine to enhance the invasiveness of the most widespread daisy in Chile, Leontodon saxatilis. Scientific Reports 7, 1546.

Mauricio R. 1998. Costs of resistance to natural enemies in field populations of the annual plant Arabidopsis thaliana. The American Naturalist 151, 20–28.

Miehe-Steier A, Roscher C, Reichelt M, Gershenzon J, Unsicker SB. 2015. Light and nutrient dependent responses in secondary metabolites of Plantago lanceolata offspring are due to phenotypic plasticity in experimental grasslands. PLoS One 10, e0136073.

Moles AT, Ackerly DD, Tweddle JC, Dickie JB, Smith R, Leishman MR, Mayfield MM, Pitman A, Wood JT, Westoby M. 2007. Global patterns in seed size. Global Ecology and Biogeography 16, 109–116.

Moles AT, Wallis IR, Foley WJ, Warton DI, Stegen JC, Bisigato AJ, Cella-Pizarro L, Clark CJ, Cohen PS, Cornwell WK, Edwards W, Ejrnaes R, Gonzales-Ojeda T, Graae BJ, Hay G, Lumbwe FC, Magana-Rodriguez B, Moore BD, Peri PL, Poulsen JR, Veldtman R, von Zeipel H, Andrew NR, Boulter SL, Borer ET, Campon FF, Coll M, Farji-Brener AG, De Gabriel J, Jurado E, Kyhn LA, Low B, Mulder CPH, Reardon-Smith K, Rodriguez-Velazquez J, Seabloom EW, Vesk PA, van Cauter A, Waldram MS, Zheng Z, Blendinger PG, Enquist BJ, Facelli JM, Knight T, Majer JD, Martinez-Ramos M, McQuillan P, Prior LD. 2011. Putting plant resistance traits on the map: a test of the idea that plants are better defended at lower latitudes. New Phytologist 191, 777–788.

Montague JL, Barrett SCH, Eckert CG. 2008. Re-establishment of clinal variation in flowering time among introduced populations of purple loosestrife (Lythrum salicaria, Lythraceae). J Evol Biol 21, 234–245.

Montaut S, Bleeker RS. 2011. Cardamine sp. – a review on its chemical and biological profiles. Chemistry & Biodiversity 8, 955–975.

Moyes CL, Collin HA, Britton G, Raybould AF. 2000. Glucosinolates and differential herbivory in wild populations of Brassica oleracea. Journal of Chemical Ecology 26, 2625–2641.

Murren CJ, Auld JR, Callahan H, Ghalambor CK, Handelsman CA, Heskel MA, Kingsolver JG, Maclean HJ, Masel J, Maughan H, Pfennig DW, Relyea RA, Seiter S, Snell-Rood E, Steiner UK, Schlichting CD. 2015. Constraints on the evolution of phenotypic plasticity: limits and costs of phenotype and plasticity. Heredity 115, 293–301.

Nahum S, Inbar M, Ne’eman G, Ben-Shlomo R. 2008. Phenotypic plasticity and gene diversity in Pistacia lentiscus L. along environmental gradients in Israel. Tree Genetics & Genomes 4, 777.

Nakagawa S, Cuthill IC. 2007. Effect size, confidence interval and statistical significance: a practical guide for biologists. Biological Reviews 82, 591–605.

Oksanen J, Blanchet FG, Friendly M, Kindt R, Legendre P, McGlinn D, Minchin PR, O’Hara RB, Simpson GL, Solymos P, Stevens MHH, Szoecs E, Wagner H. 2017. Vegan: community ecology package.

Oliva G, Martínez A, Collantes M, Dubcovsky J. 1993. Phenotypic plasticity and contrasting habitat colonization in Festuca pallescens. Canadian Journal of Botany 71, 970–977.

Pankoke H, Buschmann T, Müller C. 2013. Role of plant β-glucosidases in the dual defense system of iridoid glycosides and their hydrolyzing enzymes in Plantago lanceolata and Plantago major. Phytochemistry 94, 99–107.

Pellissier L, Fiedler K, Ndribe C, Dubuis A, Pradervand J-N, Guisan A, Rasmann S. 2012. Shifts in species richness, herbivore specialization, and plant resistance along elevation gradients. Ecology and Evolution 2, 1818–1825.

Pellissier L, Roger A, Bilat J, Rasmann S. 2014. High elevation Plantago lanceolata plants are less resistant to herbivory than their low elevation conspecifics: is it just temperature? Ecography 37, 950–959.

Pennings SC, Siska EL, Bertness MD. 2001. Latitudinal differences in plant palatability in Atlantic coast salt marshes. Ecology 82, 1344–1359.

Pilon J, Santamarìa L, Hootsmans M, van Vierssen W. 2003. Latitudinal variation in life-cycle characteristics of Potamogeton pectinatus l.: vegetative growth and asexual reproduction. Plant Ecology 165, 247–262.

Poorter H, Garnier E. 2007. Ecological significance of inherent variation in relative growth rate and its components. In: Pugnaire FI, Valladares F, eds. Functional Plant Ecology: Boca Raton: CRC Press, 67–100.

Price TD, Qvarnström A, Irwin DE. 2003. The role of phenotypic plasticity in driving genetic evolution. Proceedings of the Royal Society B: Biological Sciences 270, 1433–1440.

R Development Core Team. 2017. R: A language and environment for statistical computing. Vienna, Austria: R Foundation for Statistical Computing.

Rasmann S, Agrawal AA. 2011. Latitudinal patterns in plant defense: evolution of cardenolides, their toxicity and induction following herbivory. Ecology Letters 14, 476–483.

Rasmann S, Pellissier L, Defossez E, Jactel H, Kunstler G. 2014. Climate-driven change in plant–insect interactions along elevation gradients. Functional Ecology 28, 46–54.

Raybould A, Moyes C. 2001. The ecological genetics of aliphatic glucosinolates. Heredity 87, 383.

Ren H-X, Wang Z-L, Chen X, Zhu Y-L. 1999. Antioxidative responses to different altitudes in Plantago major. Environmental and Experimental Botany 42, 51–59.

Rodman JE, Louda SM. 1984. Phenology of glucosinolate concentrations in roots, stems and leaves of Cardamine cordifolia. Biochemical Systematics and Ecology 12, 37–46.

Rønsted N, Göbel E, Franzyk H, Jensen SR, Olsen CE. 2000. Chemotaxonomy of Plantago. Iridoid glucosides and caffeoyl phenylethanoid glycosides. Phytochemistry 55, 337–348.

Rücker B, Röbbelen G. 1994. Inheritance of total and individual glucosinolate contents in seeds of winter oilseed rape (Brassica napus L.). Plant Breeding 113, 206–216.

Savolainen O, Pyhäjärvi T, Knürr T. 2007. Gene flow and local adaptation in trees. Annual Review of Ecology, Evolution, and Systematics 38, 595–619.

Scheidel U, Bruelheide H. 2004. The impact of altitude and simulated herbivory on the growth and carbohydrate storage of Petasites albus. Plant Biology 6, 740–745.

Scheiner SM. 1993. Genetics and evolution of phenotypic plasticity. Annual Review of Ecology and Systematics 24, 35–68.

Schemske DW, Mittelbach GG, Cornell HV, Sobel JM, Roy K. 2009. Is there a latitudinal gradient in the importance of biotic interactions? Annual Review of Ecology Evolution and Systematics 40, 245–269.

Schlichting CD, Pigliucci M. 1998. Phenotypic evolution: a reaction norm perspective. Sunderland Sinauer Associates Incorporated.

Siemens DH, Mitchell-Olds T. 1998. Evolution of pest-induced defenses in Brassica plants: tests of theory. Ecology 79, 632–646.

Siska EL, Pennings SC, Buck TL, Hanisak MD. 2002. Latitudinal variation in palatability of salt-marsh plants: Which traits are responsible? Ecology 83, 3369–3381.

Spencer WE, Teeri J, Wetzel RG. 1994. Acclimation of photosynthetic phenotype to environmental heterogeneity. Ecology 75, 301–314.

Squeo FA, Rada F, Azocar A, Goldstein G. 1991. Freezing tolerance and avoidance in high tropical Andean plants: Is it equally represented in species with different plant height? Oecologia 86, 378–382.

Sultan SE. 1987. Evolutionary implications of phenotypic plasticity in plants. In: Hecht MK, Wallace B, Prance GT, eds. Evolutionary Biology, Vol. 21. Boston, MA: Springer US, 127–178.

Sultan SE. 2003. Phenotypic plasticity in plants: a case study in ecological development. Evolution & Development 5, 25–33.

Sundqvist MK, Sanders NJ, Wardle DA. 2013. Community and ecosystem responses to elevational gradients: processes, mechanisms, and insights for global change. Annual Review of Ecology, Evolution, and Systematics 44, 261–280.

Torchiano M. 2017. effsize: efficient effect size computation.

Traw MB. 2002. Is induction response negatively correlated with constitutive resistance in black mustard? Evolution 56, 2196–2205.

Van Dijk H, Wolff K, De Vries A. 1988. Genetic variability in Plantago species in relation to their ecology. Theoretical and Applied Genetics 75, 518–528.

Van Tienderen P. 1989. Measuring selection on quantitative characters: a discussion of some problems and merits of the quantitative genetic approach, applied to plant population. Grassland Species Research Group publication, 91–98.

Wagner MR, Mitchell-Olds T. 2018. Plasticity of plant defense and its evolutionary implications in wild populations of Boechera stricta. Evolution 72, 1034–1049.

Warwick SL, Briggs D. 1980. The genecology of lawn weeds. V. The adaptive significance of different growth habit in lawn and roadside populations of Plantago major L. New Phytologist 85, 289–300.

Woods EC, Hastings AP, Turley NE, Heard SB, Agrawal AA. 2011. Adaptive geographical clines in the growth and defense of a native plant. Ecological Monographs 82, 149–168.

Zehnder CB, Stodola KW, Joyce BL, Egetter D, Cooper RJ, Hunter MD. 2009. Elevational and seasonal variation in the foliar quality and arthropod community of Acer pensylvanicum. Environmental Entomology 38, 1161–1167.

Züst T, Agrawal AA. 2017. Trade-offs between plant growth and defense against insect herbivory: An emerging mechanistic synthesis. Annual Review of Plant Biology 68, 513–534.

Züst T, Rasmann S, Agrawal AA. 2015. Growth–defense tradeoffs for two major anti-herbivore traits of the common milkweed Asclepias syriaca. Oikos 124, 1404–1415.

